# Repulsive Guidance Molecule Acts in Axon Branching in *Caenorhabditis elegans*

**DOI:** 10.1101/301275

**Authors:** Kaname Tsutsui, Hon-Song Kim, Yukihiko Kubota, Yukimasa Shibata, Chenxi Tian, Jun Liu, Kiyoji Nishiwaki

## Abstract

Repulsive guidance molecules (RGMs) are evolutionarily conserved proteins implicated in repulsive axon guidance. Here we report the function of the *Caenorhabditis elegans* ortholog DRAG-1 in axon branching. The axons of hermaphrodite-specific neurons (HSNs) branch at the region abutting the vulval muscles and innervate these muscles to control egg laying. The *drag*-*1* mutants exhibited defects in HSN axon branching in addition to a small body size and egg laying–defective phenotype. DRAG-1 expression in the hypodermal cells was required for the branching of these axons. The C-terminal glycosylphosphatidylinositol anchor of DRAG-1 was important for its function. Genetic analyses suggested that the membrane receptor UNC-40, but neither SMA-1/β_H_-spectrin nor SMA-5/MAP kinase 7, acts in the same pathway with DRAG-1 in HSN branching. We propose that DRAG-1 expressed in the hypodermis signals via the UNC-40 receptor expressed in HSNs to elicit branching activity of HSN axons.

## INTRODUCTION

Axon branching is a fundamental process for proper axon projection to target tissues and for the formation of correct synapses, both of which are important components in the development of functional neural wiring. Axon branching begins with the formation of an actin-rich filopodium from the existing axon followed by extension of microtubules along the actin filaments (Sainath and Gallo 2015). Formation of filopodia and subsequent neurite extension involve various regulators for actin and microtubule polymerization and bundling. The location and the polarity of axon branching should be dictated by extracellular cues, along with cytoskeletal activities. Axon guidance molecules are involved in this process.

Netrin-1 is a secreted guidance molecule that induces local filopodial protrusions in the axon shaft, which give rise to branches in the cortical neurons. In contrast, SEMA3A represses cortical axon branching (Dent *et al.* 2004). Ephrins are membrane-bound molecules that abolish branching of thalamic axons (Mann et al., 1998). In addition to these well-known guidance molecules, repulsive guidance molecules (RGMs) also repress axon branching in cortical neurons and mossy fibers of the hippocampus (Yoshida *et al.* 2008; Shibata *et al.* 2013). In vertebrates, RGMs are glycosylphosphatidylinositol (GPI)-linked membrane proteins that constitute a family with four members: RGMa, RGMb (DRAGON), RGMc (hemojuvelin), and RGMd (Camus and Lambert 2007). RGMa was first discovered as an axon guidance cue that has a repulsive activity to retinal axons. RGMa is expressed in the embryonic tectum in an anterior-to-posterior concentration gradient and functions during the development of the retinotectal projection (Monnier *et al.* 2002; Matsunaga *et al.* 2006).

RGMs function in axon guidance and neuronal survival by binding to the membrane receptor neogenin (Matsunaga and Chedotal 2004; Rajagopalan *et al.* 2004). RGMs also bind bone morphogenetic proteins (BMPs) in the regulation of iron homeostasis and endochondral bone development (Wang *et al.* 2005; Babitt *et al.* 2006; Zhou *et al.* 2010). Although the function of RGMs in axon guidance as a result of growth cone repulsion has been well studied, their role in axon branching is still elusive. Because of the lethality of knock-out mice and the functional redundancy of RGM proteins, the functions of RGMs have been mostly analyzed using *in vitro* culture systems (Siebold *et al.* 2017).

The nematode *C. elegans* has a single ortholog of RGM, DRAG-1. Loss-of-function mutations in *drag*-*1* result in a small body size as well as genetic suppression of the coelomocyte loss phenotype of *sma*-*9* mutants (Tian *et al.* 2010). The *drag*-*1* function in the hypodermal cells is required for the control of body size and its function in the mesodermal M cell lineage controls mesodermal differentiation (Tian *et al.* 2010; Tian *et al.* 2013). In the present study, we showed that *drag*-*1* functions in the formation of branches of hermaphrodite-specific neurons (HSNs), which regulate egg laying by promoting the contraction of egg-laying muscles. We found that small body size mutants—*unc*-*40*, *sma*-*1*, *sma*-*5*, and *sma*-*8*—also exhibited HSN axon branching defects. Genetic analyses suggested that *unc*-*40* acts in the same pathway with *drag*-*1*, whereas *sma*-*1* and *sma*-*5* do not. Because DRAG-1 binds the receptor UNC-40 (Tian *et al.* 2013) and UNC-40 is expressed in HSNs (Tang and Wadsworth 2014), it is possible that DRAG-1 acts on UNC-40 to induce axon branching of HSNs. Our findings indicate that DRAG-1 promotes axon branching in contrast to other RGMs, which inhibit axon branching in cortical neurons and mossy fibers of the hippocampus (Yoshida *et al.* 2008; Shibata *et al.* 2013).

## MATERIALS AND METHODS

### Strains and culture conditions

Culture and handling of *C. elegans* were as described (Brenner 1974). The following strains were used: N2 (wild type, WT), *drag*-*1(tk81)* (this work), *drag*-*1(tm3773)* (National Bioresource Project), *unc*-*119(e2498)* (Maduro and Pilgrim 1995), *sma*-*1(e30)*, *sma*-*2(e502)*, *sma*-*3(wk28)*, *sma*-*4(e729)*, *sma*-*5(n678)*, *sma*-*6(wk7)*, *sma*-*8(e2111)*, *sma*-*9(wk55)*, *unc*-*40(e271)* (Brenner 1974; Savage-Dunn *et al.* 2003; Watanabe *et al.* 2005). HSNs were visualized using an integrated transgene *kyIs262[unc*-*86p::myrGFP; odr*-*1::RFP]* (Adler *et al.* 2006). *drag*-*1(tk81)* was isolated by the trimethylpsoralen and UV irradiation method (Kubota *et al.* 2004).

### Plasmid construction

*drag*-*1p::drag*-*1::gfp::GPI*, *drag*-*1p::drag*-*1::gfp::ΔGPI*, and *drag*-*1p::drag*-*1::gfp::lin*-*12TM* correspond to pJKL849, pCX192, and pCX194, respectively (Tian *et al.* 2010). To produce *drag*-*1p::venus*, the *drag*-*1* promoter region was PCR amplified from genomic DNA using primers 5′-TAGCCTGCAGGTTTCCGAAGACAGGGGAACATGGAA-3′ and 5′-GTTCGTCGACACTCTGTCAAGTCTTCTCATCTCACG-3′, digested with *Pst*I and *Sal*I, and cloned into the *Pst*I and *Sal*I sites of pPD95.75. To produce *drag*-*1p::drag*-*1::venus*, the *drag*-*1* coding region was PCR amplified from genomic DNA with primers 5′-GTCAGTCGACATGTCAATAGTCTATCTCG-3′ and 5′-CATGGGTACCAAGCATAACAATGATAAAAGAGC-3′, digested with *Sal*I and *Kpn*I, and cloned into the *Sal*I and *Kpn*I sites of *drag*-*1p::venus.* To produce *drag*-*1p::drag*-*1*, *drag*-*1p::drag*-*1::venus* was PCR amplified with primers 5′-GATCGCTAGCCTTGTCTGGTGTCAAAAATAATAGG-3′ and 5′ - TCGCTAGCTCAGCATAACAATGATAAAAGAGCAAAA-3′, digested with *Nhe*I, and self-ligated. To produce *drag*-*1p::drag*-*1::venus::GPI*, the Venus coding region was PCR amplified from pPD95.75 with primers 5′-GCATGGGCCCAGGGTACCGGTAGAAAAAATGAGT-3′ and 5′-GCATGGGCCCTTTGTATAGTTCATCCATGCCAAG-3′, digested with *Apa*I, and ligated into *drag*-*1p::drag*-*1::gfp::GPI* (pJKL849) in which the green fluorescent protein (GFP) coding region had been deleted by *Apa*I digestion. *rol*-*6*, *myo*-*2*, and *elt*-*2* promoter regions were PCR amplified with primers 5′-CAGTGCATGCCGAGAAGAGTCCGGTGTGAA-3′ and 5′-CAGTGTCGACCTGGAAATTTTCAGTTAGATCTAAAG-3′, 5′-CAGTGCATGCGTGAGCAAGTGTGCGGCATC-3′ and 5′-CAGTGTCGACTTCTGTGTCTGACGATCGAGGG-3′, and 5′-CAGTCCTGCAGGGTGACCGCTCAAAATAAAAGG-3′ and 5′-CAGTCTCGAGTCTATAATCTATTTTCTAGTTTCTA-3′, respectively. These PCR fragments were digested with *Pst*I and *Sal*I and ligated with *drag*-*1p::drag*-*1* in which the *drag*-*1* promoter region was deleted by *Pst*I and *Sal*I digestion to produce *rol*-*6p::drag*-*1*, *myo*-*2p::drag*-*1*, and *elt*-*2p::drag*-*1*.

### Production of transgenic animals

We injected DNA mixtures into the gonads of *unc*-*119(e2498)*, *drag*-*1(tk81); unc*-*119(e2498); kyIs262 or drag*-*1(tm3773); unc*-*119(e2498); kyIs262* adult hermaphrodites (Mello *et al.* 1991). For transgenic rescue experiments, test plasmids were injected at 10–20 ng/μ1 with 160–170 ng/μl of pBSII KS(–) and 20 ng/μl of *unc*-*119*^+^ plasmid pDP#MM016B (Maduro and Pilgrim 1995). For immunohistochemistry, *drag*-*1p::drag*-*1* and *drag*-*1p::drag*-*1::venus::GPI* plasmids were injected at 150 ng/μl with 30 ng/μl of pBSII KS(–) and 20 ng/μl pDP#MM016B.

### Production of antibodies

The RNA sample extracted from wild-type worms was treated with SuperScript III reverse transcriptase (Invitrogen) using a primer 5′-TCAGCATAACAATGATAAAAGAGC-3′ designed to anneal at the 3′-end of the coding region, and single-strand cDNA was produced. The double-strand cDNA was amplified by PCR using a primer designed to anneal at the SL1 splice leader sequence 5′-GGTTTAATTACCCAAGTTTGAG-3′ and the 3′-end primer. The region coding for DRAG-1 peptide from I131 to E368 was amplified using primers 5′-GTCACATATGATAATGTTCAATGGCTCCGTGC-3′ and 5′-GTCACTCGAGTTCTTTCTGGAACCGAGCATG-3′, digested with *Nde*I and *Xho*I, and ligated into the pET-19b vector using the *Nde*I and *Xho*I sites. The resulting antigenic peptide of DRAG-1 was expressed as a histidine-tagged fusion protein in *Escherichia coli* and was used to immunize rabbits. The generated antibody was affinity purified.

### Immunohistochemistry

Immunohistochemistry was performed as described (Kim *et al.* 2011). The DRAG-1 antibody was used as the primary antibody at 4 μg/ml. Alexa 594–labeled donkey anti–rabbit IgG (Life Technologies) was used as the secondary antibody at a dilution of 1:500.

### Microscopy

Nomarski and fluorescence microscopy was performed using an Axioplan 2 microscope equipped with Axiocam CCD camera (Zeiss). Confocal laser scanning microscopy was conducted with LSM5 (Zeiss) equipped with a C-apochromat 63× (water immersion; NA, 1.2) lens controlled by PASCAL version 3.2 SP2 software.

### Data availability

Strains and plasmids are available upon request. The authors state that all data necessary for confirming the conclusions presented in the article are represented fully within the article.

## RESULTS

### *drag*-*1* mutants are defective in axon branching of HSNs

*drag*-*1* encodes the sole *C. elegans* ortholog of the RGM family of proteins. We isolated a deletion allele of *drag*-*1*, *tk81.* The *drag*-*1(tk81)* animals had a smaller body size compared to wild type and showed a partial egg laying–defective phenotype. The *tm3773* mutants showed a similar phenotype (Figure 1, A-C). Egg laying is mainly regulated by the HSNs, which innervate vulval muscles (White *et al.* 1986). Therefore, we examined the morphology of the HSNs using *unc*-*83p::myrGFP* as a transgenic marker. The cell bodies of the bilateral HSNs are positioned slightly posterior to the vulva. They extend a single axon toward the ventral nerve cord. After reaching the nerve cord, the axon is redirected dorsally and anteriorly and reaches the lateral position of the vulva, where it turns again, this time in a ventral and anterior direction to fasciculate with the ventral nerve cord (Garriga *et al.* 1993). Although the axon usually sprouts one or two branches at the vulva in the wild type, the number of axons with branches was significantly reduced in the *drag*-*1* mutants (Figure 2, A and B).

**Figure 1.**
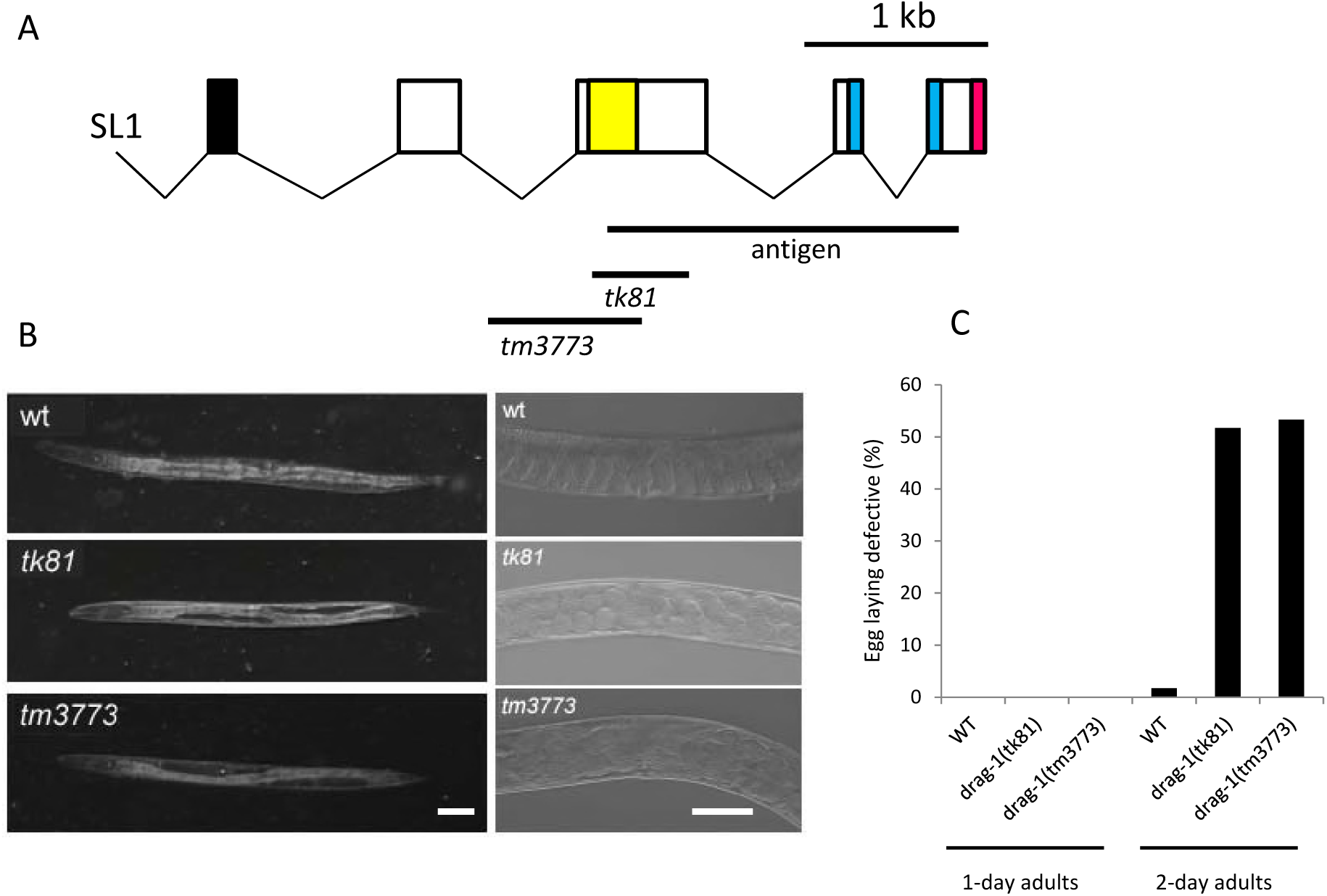
Gene structure and mutant phenotype of *drag*-*1* animals. (A) Structure of *drag*-*1* and mutation sites of the *tk81* and *tm3773* alleles. The exon and intron regions were determined by sequencing cDNA generated from isolated *drag*-*1* mRNA. SL1, splice leader sequence 1. Black, yellow, blue, and magenta boxes indicate N-terminal signal peptide, partial von Willebrand factor type D domain, hydrophobic region, and C-terminal GPI-anchor signal sequence, respectively (Tian *et al.* 2010). Bars depict the region of the cDNA used for expressing the antigenic peptide for producing antibodies and the mutation sites. *tk81* is a 494-bp deletion within exon 3, which is expected to produce a truncated polypeptide that is missing the C-terminal 278 amino acids. *tm3773* is an 892-bp deletion spanning from intron 2 to exon 3 (WormBase). (B) Body length and egg-laying phenotype of *drag*-*1* mutants. Left panels: Body length of young adult hermaphrodites. *tk81* and *tm3773* mutants had shorter bodies compared with wild type. Right panels: The egg laying–defective phenotype was assessed by the accumulation of fertilized eggs within the uterus in 2-day-old adult hermaphrodites. *tk81* and *tm3773* mutants often accumulate eggs, but wild-type animals do not. Anterior is to the left. Bar: 50 μm. (C) Percentages of egg laying–defective animals in wild-type and *drag*-*1* mutant animals. Data for 1-day-old and 2-day-old adults are shown. N = 60 for all experiments.

**Figure 2.**
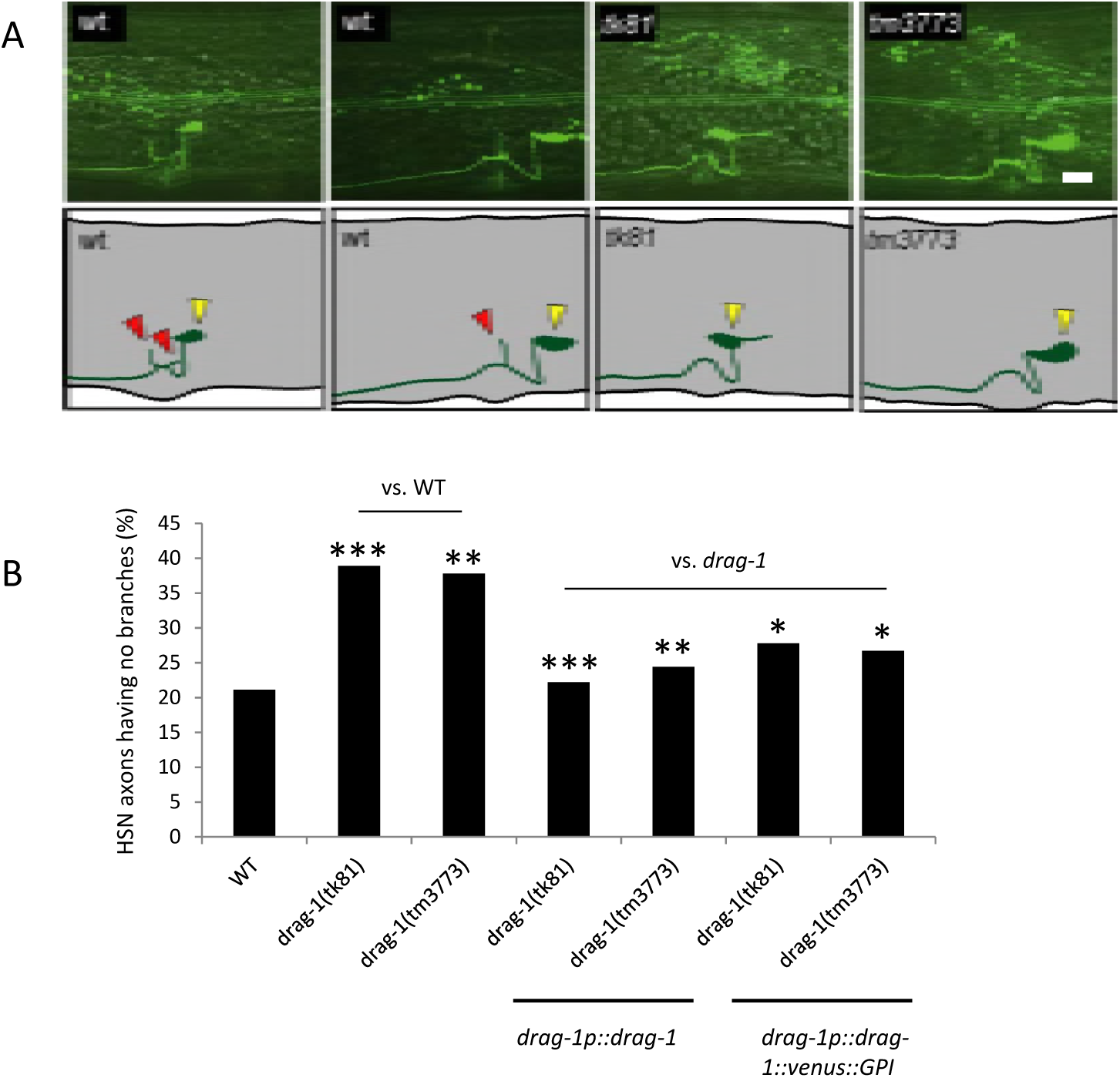
HSN branching phenotypes. (A) HSN axon branching. Upper panels: Confocal micrographs of HSNs in wild-type and *drag*-*1* mutant young-adult hermaphrodites with the *unc*-*86p::myrGFP* transgene. Lower panels: Schematic representations of HSN morphology. Yellow and red arrowheads depict the HSN cell body and axonal branches, respectively. Anterior is to the left, dorsal top. Bar: 10 μm. (B) Transgenic rescue of *drag*-*1* mutants. Percentages of HSNs having no branches are shown. Branching phenotypes were scored using fluorescence microscopy. *drag*-*1(tk81)* and *drag*-*1(tm3773)* mutants were compared with those transgenic for *drag*-*1p::drag*-*1* and *drag*-*1p::drag*-*1::venus::GPI.* Significant differences were determined by Fisher’s exact test relative to WT for *drag*-*1(tk81)* and *drag*-*1(tm3773)* and relative to each *drag*-*1* mutant for the transgenic strains. ^***^P < 0.001, ^**^P < 0.01, ^*^P < 0.05. N = 180 for all experiments.

### GPI anchoring of DRAG-1 is important for HSN branching

To examine whether the branching defect is caused by loss of *drag*-*1* function, we introduced a plasmid containing a fragment of the wild-type gene (*drag*-*1p::drag*-*1*) into the *drag*-*1* mutants. The plasmid fully rescued the branching defect (Figure 2B), confirming the function of DRAG-1 in HSN axon branching. Because DRAG-1 is thought to be modified by a GPI-anchor, we placed the GFP or Venus coding region right upstream of the putative cleavage site of the C-terminal pro-peptide sequence (*drag*-*1p::drag*-*1::gfp::GPI* or *drag*-*1p::drag*-*1::venus::GPI*) to retain the GPI-anchor signal intact. These constructs rescued the branching defect of the mutants (Figure 2B, Figure 3, A and B). We examined whether the GPI anchoring is important for DRAG-1 to act in axon branching. A construct with the deleted C-terminal GPI-anchor signal sequence (*drag*-*1p::drag*-*1::gfp::ΔGPI*) failed to rescue the branching defects of *drag*-*1* mutants (Figure 3, A and B). We also examined a construct in which the C-terminal GPI-anchor signal was replaced by the transmembrane domain of the LIN-12 receptor (*drag*-*1p::drag*-*1::gfp::lin*-*12TM*). This chimeric protein, which potentially localizes to the plasma membrane, failed to rescue the mutant defects (Figure 3, A and B). These results suggested that DRAG-1 should be anchored to the plasma membrane by the GPI anchor and that the anchoring by the LIN-12 transmembrane domain can abrogate DRAG-1 function.

**Figure 3.**
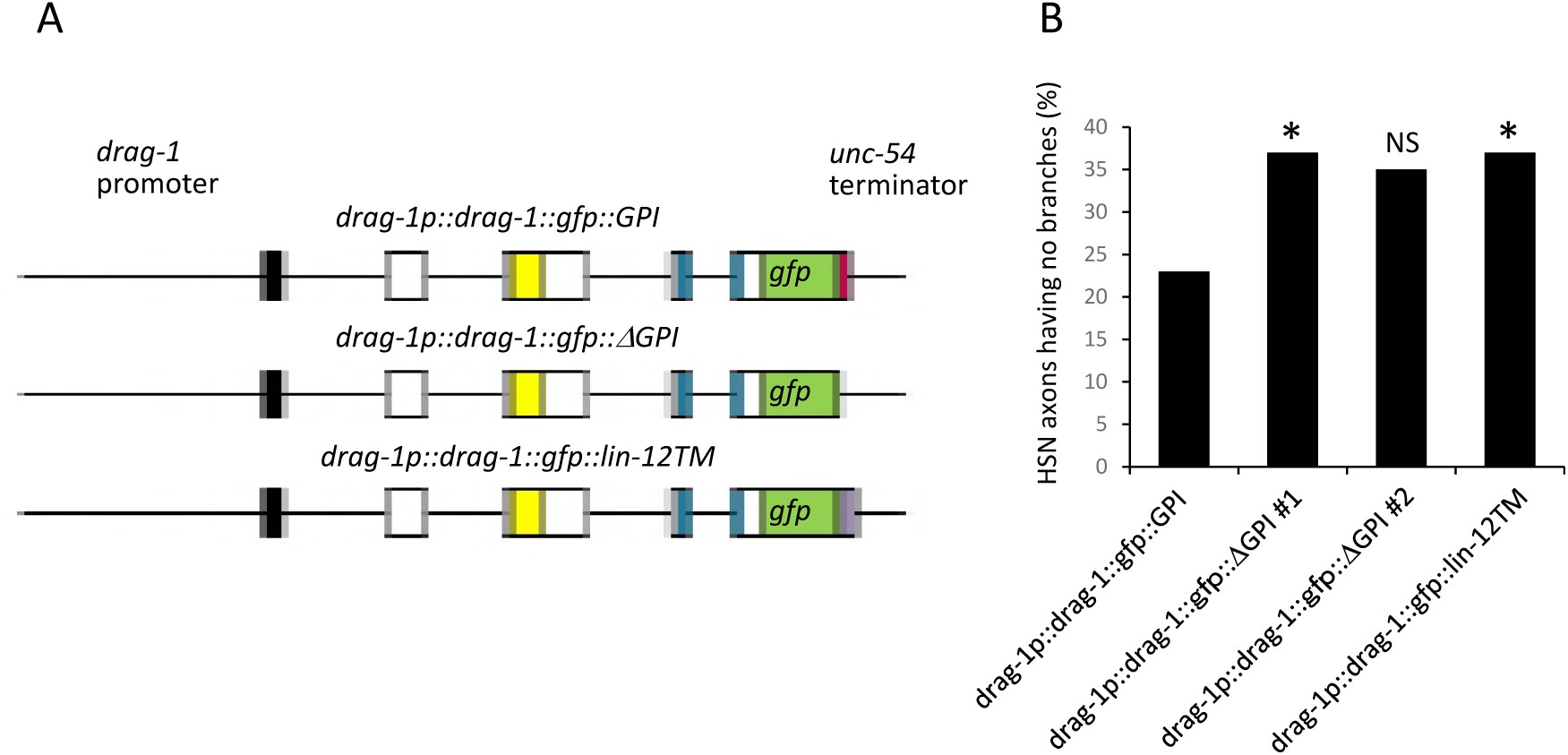
Rescue experiments of *drag*-*1* mutants with modified DRAG-1 proteins. (A) Schematic presentation of the GFP fusion constructs. The GFP coding sequence was inserted between amino acid (aa) 395 and 396 of the *drag*-*1* coding region, just prior to the cleavage site of the C-terminal pro-peptide for *drag*-*1p::drag*-*1::gfp::GPI.* The C-terminal GPI-anchor signal (aa387–408) was deleted from *drag*-*1p::drag*-*1::gfp::GPI* for *drag*-*1p::drag*-*1::gfp::ΔGPI*. The *lin*-*12* transmembrane domain (aa 907–934) (shown in purple) was connected with *drag*-*1p::drag*-*1::gfp:: ΔGPI* for *drag*-*1p::drag*-*1::gfp::lin*-*12TM*(Tian et al. 2010). (B) Percentages of HSNs having no branches are shown for the *drag*-*1(tk81)* mutants that also expressed the constructs in (A). Significant differences were determined by Fisher’s exact test relative to *drag*-*1p::drag*-*1::gfp::GPI.* ^*^P < 0.05. NS, not significant. N = 107, 105, 91, and 109 for *drag*-*1p::drag*-*1::gfp::GPI*, *drag*-*1p::drag*-*1::gfp::ΔGPI #1*, *drag*-*1p::drag*-*1::gfp::ΔGPI #2*, and *drag*-*1p::drag*-*1::gfp::lin*-*12TM*, respectively. The *#1* and *#2* refer to two independently isolated transgenic lines.

### Expression of DRAG-1

We examined the expression of *drag*-*1* using a transcriptional *drag*-*1p::venus* reporter construct. Venus expression was detected in the pharynx, intestine, and hypodermis from late embryo to adult stages (Figure 4A) as observed by GFP fusion (Tian *et al.* 2010). Unlike the transcriptional reporter described above, we could only detect Venus expression in the pharynx using the functional translational fusion construct *drag*-*1p::drag*-*1::venus::GPI* (Figure 4B). We raised polyclonal antibodies against a DRAG-1 peptide corresponding to amino acids 130–368 (Figure 1A). Immunostaining experiments indicated that the antibodies detected no signals in non-transgenic wildtype animals. However, they detected signals in those animals transgenic for *drag*-*1p::drag*-*1* or *drag*-*1p::drag*-*1::venus::GPI* in a similar pattern. The signals were detected in the pharynx, intestinal cells, hypodermal seam cells, and in the ventral hypodermal cells except in the vulval region (Figure 4D). Thus, it is likely that the level of expression of endogenous DRAG-1 is low. The hypodermal signals detected by the anti-DRAG-1 antibodies appeared in a granular pattern in the cytoplasm. It is possible that DRAG-1 protein may be localized to the ER or the Golgi apparatus in these cells. However, we cannot rule out the possibility that the localization pattern is due to overexpression of *drag*-*1*.

**Figure 4.**
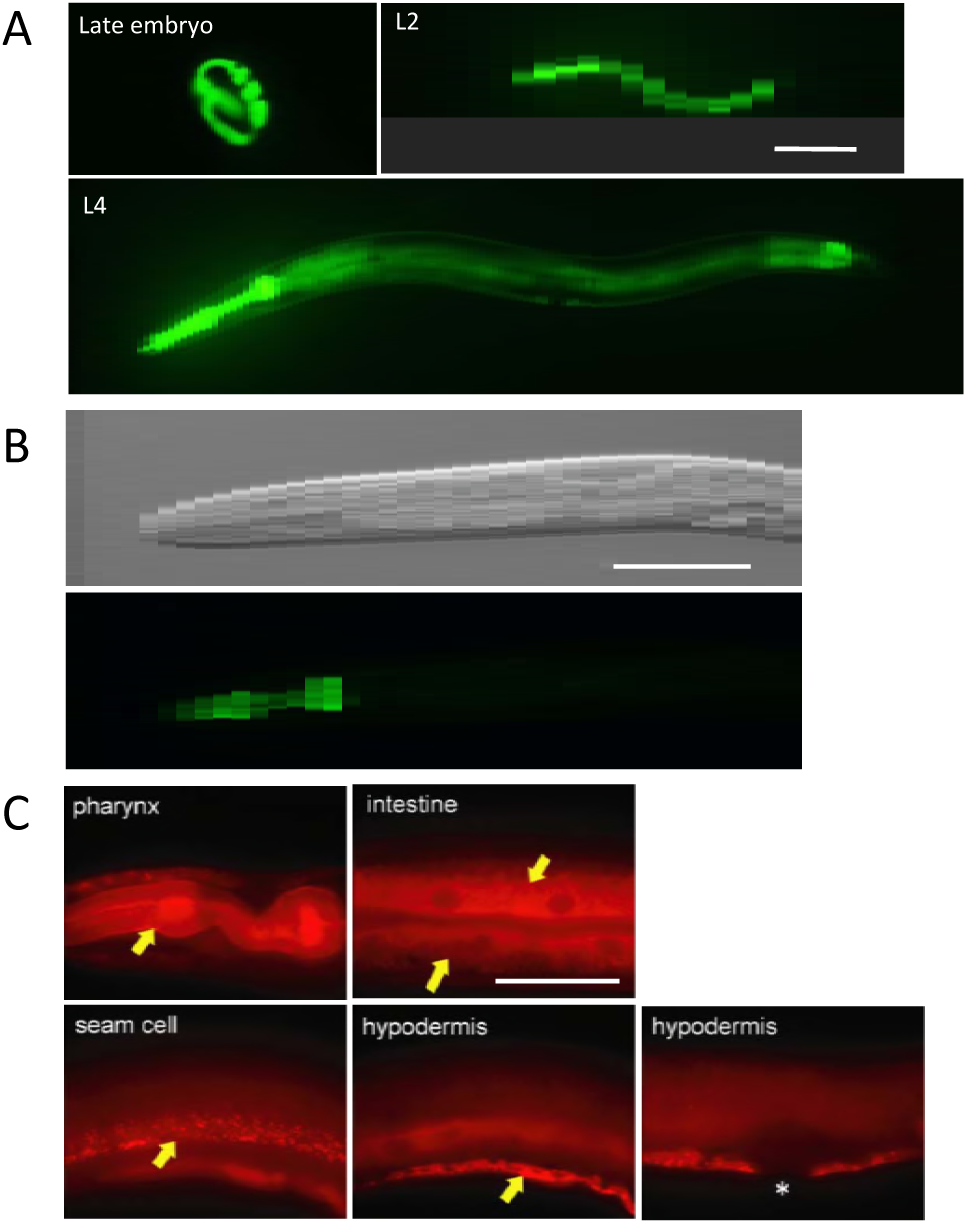
*drag*-*1* expression. (A) *drag*-*1p::venus* expression. Expression was detected from late embryos to the adult stage in the pharynx, intestine, and hypodermis. Bar: 100 μm. (B) Expression of *drag*-*1p::drag*-*1::venus::GPI.* The *drag*-*1p::drag*-*1::venus::GPI* plasmid was injected into *unc*-*119(e2498)* animals at 150 ng/μl with 30 ng/μl of pBSII KS(–) and 20 ng/μl pDP#MM016B. DIC (upper) and fluorescence (lower) images of an L4 stage animal are shown. Venus expression was detected only in the pharynx. Anterior is to the left. Bar: 100 μm. (C) Immunostaining using anti-DRAG-1. L4 to young-adult animals expressing *drag*-*1p::drag*-*1::venus::GPI* were stained with anti-DRAG-1. DRAG-1 expression was detected in the pharynx, intestine, hypodermal seam cells, and ventral hypodermal cells (arrows) with the exception of the vulval hypodermis (asterisk). Bar: 50 μm.

### DRAG-1 functions in hypodermal cells for axon branching

To determine the tissues in which DRAG-1 expression is important for axon branching, we expressed *drag*-*1* under tissue-specific promoters. We found that hypodermal expression of DRAG-1 using the *rol*-*6* promoter (*rol*-*6p::drag*-*1*) rescued the branching defect, whereas expression in the pharyngeal muscle (*myo*-*2p::drag*-*1*) or in the intestine (*elt*-*2p::drag*-*1*) did not (Figure 5). These results indicated that DRAG-1 functions non-cell-autonomously in hypodermal cells to induce axon branching of the HSNs.

**Figure 5.**
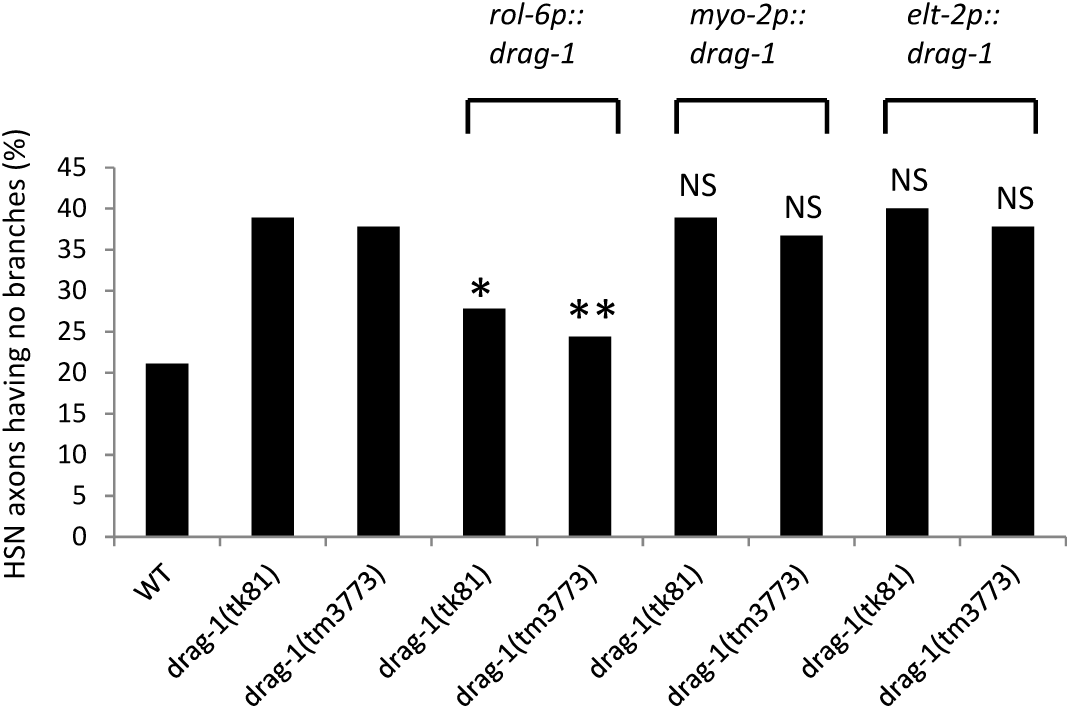
Tissue-specific rescue experiments of *drag*-*1* mutants. Percentages of HSNs having no branches are shown. *drag*-*1(tk81)* and *drag*-*1(tm3773)* mutants were compared with those transgenic for *rol*-*6p::drag*-*1*, *myo*-*2p::drag*-*1*, and *elt*-*2p::drag*-*1.* Significant differences were determined by Fisher’s exact test relative to *drag*-*1* mutants. ^**^P < 0.01, ^*^P < 0.05. NS, not significant. N = 180 for all experiments.

### *drag*-*1* acts in parallel pathways with *sma*-*1* and *sma*-*5*

Because *drag*-*1* mutants result in a small body size (Sma) phenotype, we examined whether other *sma* mutants affect HSN axon branching. Among the eight *sma* mutants examined, *sma*-*2(g502)*, *sma*-*3(wk28)*, *sma*-*4(e728)*, *sma*-*6(wk7)*, and *sma*-*9(wk55)* did not show HSN branching defects (Figure 6A). We found HSN branching defects similar to that observed in the *drag*-*1* mutants in *sma*-*1(e30)*, *sma*-*5(n678)*, and *sma*-*8(e2111)* (Figure 6B). *sma*-*1* and *sma*-*5* encode β_H_-spectrin and MAP kinase 7, respectively (McKeown *et al.* 1998; Geisler *et al.* 2016). *sma*-*8(e2111)* is a dominant mutation for which the causative gene has not yet been identified. We produced double mutants between *drag*-*1* mutants and these *sma* mutants and found that all double mutants exhibited HSN branching defects stronger than those observed in the respective single mutants (Figure 6B). Because *drag*-*1(tk81)* and *drag*-*1(tm3773)* mutants are putative null alleles, these results suggested that *sma*-*1* and *sma*-*5* act in pathways different from that of *drag*-*1* to regulate HSN branching. The relationship between genetic pathways for *drag*-*1* and *sma*-*8* is not clear because of the dominancy of the *sma*-*8(e2111)* mutation.

**Figure 6.**
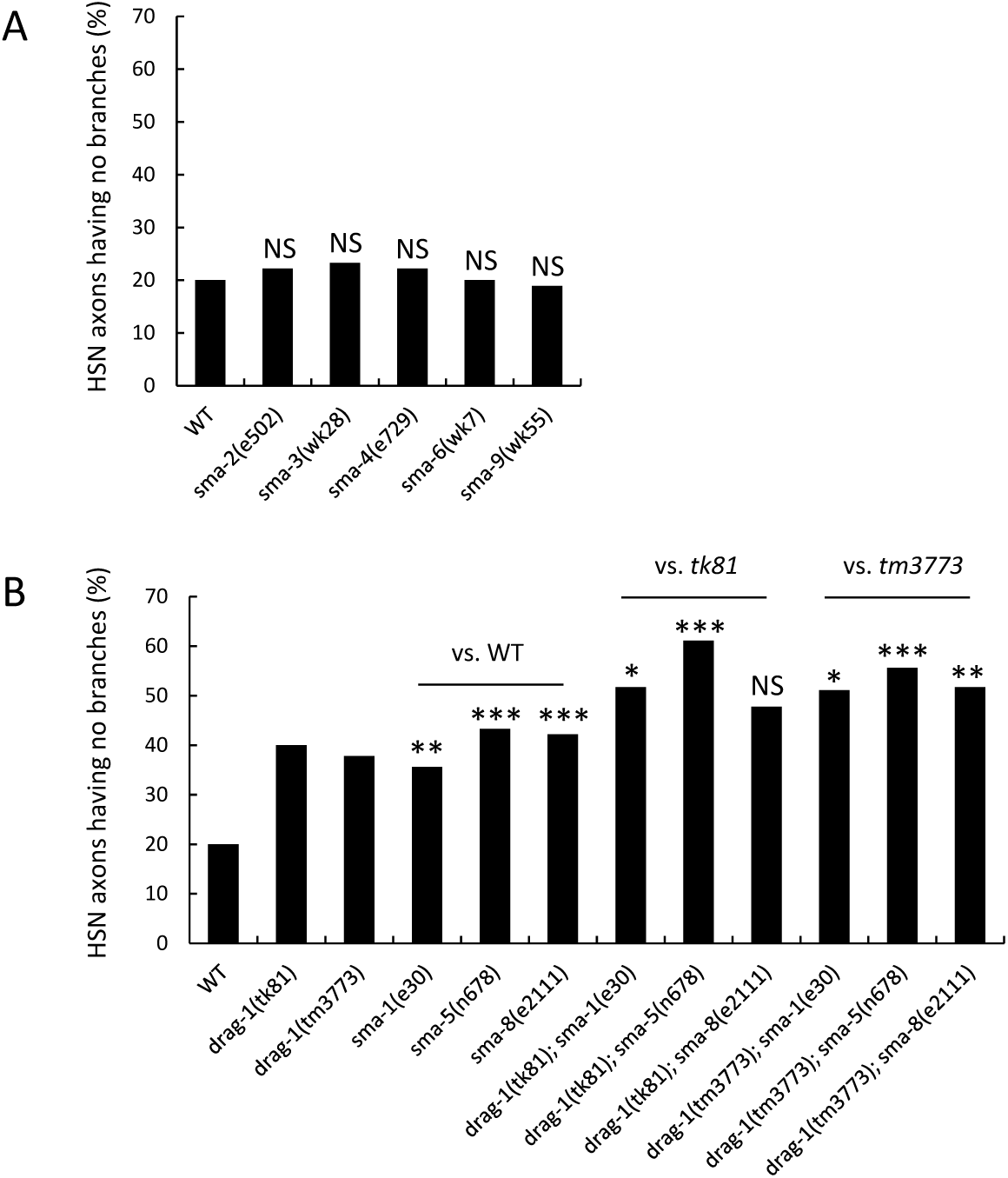
Genetic interactions between *drag*-*1* mutants and *sma* mutants. Percentages of HSNs having no branches are shown. (A) *sma*-*2(g502)*, *sma*-*3(wk28)*, *sma*-*4(e728)*, *sma*-*6(wk7)*, and *sma*-*9(wk55)* mutants were compared with wild type. (B) *drag*-*1(tk81)* and *drag*-*1(tm3773)* mutants were compared with *sma*-*1(e30)*, *sma*-*5(n678)*, and *sma*-*8(e2111)* mutants and with double mutants consisting of *drag*-*1(tk81)* or *drag*-*1(tm3773)* in combination with individual *sma* mutations. Significant differences were determined by Fisher’s exact test. ^***^P < 0.001, ^**^P < 0.01, ^*^P < 0.05. NS, not significant. N = 180 for all experiments.

### *drag*-*1* acts in the same pathway with *unc*-*40*

Neogenin is a receptor for RGMa for axonal growth cone guidance (Matsunaga and Chedotal 2004; Rajagopalan *et al.* 2004). UNC-40, a well-known receptor for UNC-6/netrin (Culotti and Merz 1998), is the sole ortholog of neogenin in *C. elegans.* Although we tried to examine HSN branching defects in *unc*-*40(e271)* null mutants, the severe axon guidance defect in these mutants—which is likely to be caused by disruption of the UNC-6-dependent guidance signaling—made it impossible to examine the branching phenotype. However, we observed branching defects in *unc*-*40(e271)/*+ heterozygotes with similar penetrance as in the *drag*-*1* mutants. This defect was not enhanced when combined with *drag*-*1* null mutants (Figure 7). Therefore, UNC-40 acts in the same genetic pathway with DRAG-1 in HSN axon branching.

**Figure 7.**
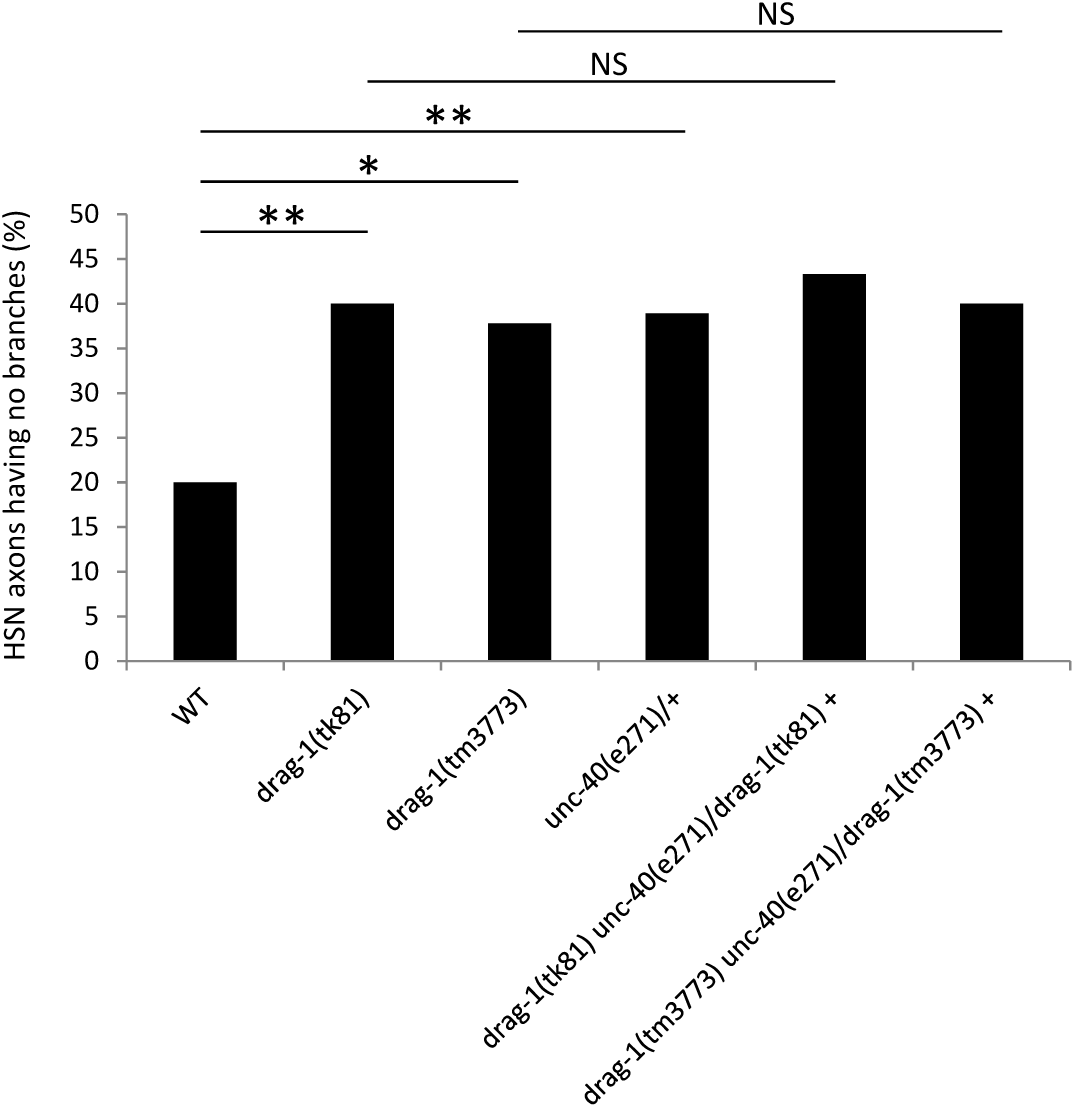
*drag*-*1* does not enhance *unc*-*40/*+ with respect to HSN branching defects. Percentages of HSNs having no branches are shown. *drag*-*1(tk81)* and *drag*-*1(tm3773)* mutants were compared with *unc*-*40(e271)/*+ heterozygotes and with *drag*-*1(tk81) unc*-*40(e271)/*+ and *drag*-*1(tm3773) unc*-*40(e271)/*+ double mutants. Significant differences were determined by Fisher’s exact test. ^**^P < 0.01, ^*^P < 0.05. NS, not significant. N = 90 for all experiments.

## DISCUSSION

### DRAG-1 acts in axon branching

In the present study, we found that DRAG-1, the sole ortholog of RGMs in *C. elegans*, acts in axon branching of the HSN, which is required for egg laying. Immunohistochemistry using a DRAG-1 antibody revealed that DRAG-1 is expressed in hypodermal seam cells and in the ventral hypodermis with the exception of the vulval epithelium. The defective axon branching of *drag*-*1* mutants was rescued by *rol*-*6p::drag*-*1.* Because the *rol*-*6* promoter drives gene expression in the hypodermis excluding the seam cells (Sassi *et al.* 2005), it is possible that DRAG-1 expression in the ventral hypodermis is important for HSN axon branching. Thus, DRAG-1 expressed in the ventral hypodermal cells is likely to induce HSN branching in a non-cell-autonomous fashion. Genetic analysis suggested that DRAG-1 acts through the receptor UNC-40, which is expressed in the HSNs. This is consistent with UNC-40 being an ortholog of vertebrate neogenin, which acts as a receptor for RGM proteins (Rajagopalan *et al.* 2004; Cole *et al.* 2007).

The branches of HSNs form during vulval morphogenesis during the fourth larval stage (Asakura *et al.* 2007). Although the branching region of the HSN may make direct contact with the vulval epithelium or hypodermal seam cells, it does not contact the ventral hypodermis. So how can DRAG-1 transduce a signal via UNC-40? We suggest two possibilities. First, DRAG-1 expressed as a GPI-anchored protein in the ventral hypodermis may be detached from the plasma membrane and diffuse to the site of branching of HSNs, where it can bind UNC-40 to elicit the branching signal. Mammalian RGMc, but not RGMa or RGMb, is cleaved by furin at a specific C-terminal site (Silvestri *et al.* 2008). Second, UNC-40 is expressed in both the cell body and in the axons of the HSNs (Tang and Wadsworth 2014). HSN axons fasciculate with the ventral nerve cord twice at regions posterior and anterior to the vulva (Garriga *et al.* 1993). Because the ventral nerve cord runs along the ventral hypodermis, it is possible that DRAG-1 expressed on the ventral hypodermal surface directly binds UNC-40 expressed in the HSN axons to transduce the signal. Also, DRAG-1 physically interacts with the extracellular domain of UNC-40 in *C. elegans* (Tian *et al.* 2013) as observed for the interaction between human RGMc and neogenin (Yang *et al.* 2008).

Because neither DRAG-1 with its GPI-anchor sequence deleted (therefore a potential secreted form) nor DRAG-1 fused with the LIN-12 transmembrane domain (therefore a potential membrane-anchored form) rescued the *drag*-*1* mutants, we cannot distinguish between these two possibilities. The latter was unexpected because the same construct significantly rescues the *drag*-*1* defect in the control of mesodermal cell differentiation (Tian *et al.* 2010). With respect to mesodermal cell differentiation, DRAG-1 and UNC-40 are expressed in the same cells to promote BMP signaling. In HSN branching, however, they are expressed in different cell types. Thus it is possible that membrane tethered DRAG-1 can act on the UNC-40 receptor cell autonomously, but not non-cell-autonomously.

### Egg laying and HSN axon branching

HSN axons form varicosities and branches in the region of the vulva within which the HSNs form synapses on egg-laying muscles. The neurotransmitter serotonin, which induces muscle contraction, activates egg laying (White *et al.* 1986; Desai *et al.* 1988; Garriga *et al.* 1993). We observed HSN branching defects in about 40% of *drag*-*1* mutants (compared to 20% in wild type), but we found that about 50% of 2-day *drag*-*1* mutant adults retained eggs, while only 2% wild-type animals did. Because HSNs also form synapses on egg-laying muscles, and secret serotonin to active egg laying, it is possible that *drag*-*1* mutants not only affect HSN axon branching, but also HSN synapse formation.

### Multiple mechanisms of HSN axon branching

Mutations in various genes result in a small body size (Sma) phenotype. Among these genes, *dbl*-*1* (BMP); *sma*-*6* (BMP type I receptor); *sma*-*2*, -*3*, and -*4* (Smads); and *sma*-*9* (BMP antagonist schnurri) are components of BMP signaling, regulating mesodermal cell fate in *C. elegans. drag*-*1* acts at the ligand-receptor level during this BMP signaling (Tian *et al.* 2010). However, none of the *sma* mutants in the BMP pathway affected HSN axon branching. Instead, we observed HSN axon branching defects in *sma*-*1(e30)* and *sma*-*5(n678)* mutants, which are not involved in the BMP signaling, similar to that observed in *drag*-*1* mutants. Therefore, the BMP signaling does not regulate HSN axon branching and DRAG-1 regulate HSN axon branching independent of the BMP signaling.

SMA-1 and SMA-5 appear to act in pathways parallel to that of DRAG-1. *sma*-*1* encodes β_H_-spectrin, which is a very large spectrin found in invertebrates such as *C. elegans* and *Drosophila* (McKeown *et al.* 1998). The submembrane skeletal network is primarily formed from α2β2 spectrin tetramers, each composed of two α-spectrin and two β-spectrin subunits (Bennett and Gilligan 1993). The spectrin network interacts with peripheral actin filaments to act in synapse function, muscle sarcomere structure, and axonal outgrowth (Hammarlund *et al.* 2000; Moorthy *et al.* 2000). Although SMA-1 function in shaping cells in the hypodermis and pharyngeal muscles has been reported (Praitis *et al.* 2005; Raharjo *et al.* 2011), its function in neuronal cells is unknown. SMA-1 may function in HSNs for branch formation by regulating actin filaments. SMA-5/MAP kinase 7 is specifically expressed in the intestine to control intestinal tube stability and body size (Geisler *et al.* 2016). Because the intestine has no direct contact with HSNs, it is possible that SMA-5 indirectly affects branching of the neuron. Because 60% of HSNs produce at least one branch in the *drag*-*1* null mutant background and 40–50% of HSNs still make branches even in *drag*-*1* and *sma*-*1* or *sma*-*5* double mutants, it is likely that multiple mechanisms operate in branch formation in HSNs.

In summary, we provide in vivo evidence that RGM proteins function to promote axon branching. Our finding is in contrast to the observation that RGM proteins suppress the branching of axons in the mammalian brain. RGMs may function in both ways depending on the tissues or the phases of organogenesis. Further research is needed to understand the precise function of RGMs in axon branching.

## ACKNOWLEDGMENTS

We thank Noriko Nakagawa and Nami Okahashi for technical assistance. Some nematode strains used in this work were provided by the Caenorhabditis Genetics Center, which is funded by the National Institutes of Health National Center for Research Resources and by Shohei Mitani through the National Bioresource Project for the nematode. This work was supported by This work was supported by NIH R01 GM103869 to J.L., and a Grant-in-Aid for Scientific Research by Ministry of Education, Culture, Sports, Science and Technology to KN.

